# Lysosomal dysfunction impairs mitochondrial quality control and predicts neurodegeneration in TBCKE

**DOI:** 10.1101/2020.05.25.114306

**Authors:** Jesus A Tintos-Hernandez, Kierstin N. Keller, Adrian Santana, Xilma R Ortiz-Gonzalez

**Author notes:** **Corresponding Author**, Xilma R Ortiz-González, MD PhD, Assistant Professor of Neurology and Pediatrics, University of Pennsylvania, The Children’s Hospital of Philadelphia, 3501 Civic Center Blvd, Philadelphia, PA 19104.

## Abstract

Biallelic variants in TBC1-domain containing kinase (*TBCK*) cause intellectual disability in children. It remains unclear how variants in *TBCK* lead to a neurodevelopmental disorder and what biological factors modulate the variability of clinical severity. Previous studies showed increased autophagosomes in patients sharing the truncating (p.R126X) Boricua homozygous *TBCK* variant, who exhibit a severe and progressive neurodegenerative phenotype. Since defects in mitophagy are linked to neurodegenerative disorders, we tested whether mitophagy and mitochondrial function are altered in *TBCK*^-/-^ fibroblasts. Our data shows significant accumulation of mitophagosomes, reduced mitochondrial respiratory capacity, and mtDNA depletion. Furthermore, mitochondrial dysfunction correlates with the severity of the neurological phenotype. Since effective mitophagy and degradation of mitophagosomes ultimately depends on successful lysosomal degradation, we also tested lysosomal function. Our data shows that lysosomal proteolytic function is significantly reduced in *TBCK*^-/-^ fibroblasts. Moreover, acidifying lysosomal nanoparticles rescue the mitochondrial respiratory defects, suggesting that impaired mitochondrial quality control secondary to lysosomal dysfunction, may play an important role in the pathogenicity of this rare neurodevelopmental disorder and predict the degree of disease progression and neurodegeneration.

## Introduction

Genetic variants causing mitochondrial and lysosomal disorders often present clinically as pediatric neurodegenerative diseases. On the broader spectrum of common neurologic disorders, defects in these critical organelles have being increasingly implicated in the pathogenesis of Parkinson’s and Huntington’s disease (Menzies *et al,* 2015; Schon & Przedborski, 2011; Wallace, 2011; Whyte *et al,* 2017). We have previously reported that biallelic mutations in the *TBCK* gene cause intellectual disability of variable severity in children, with some patients presenting with intellectual disability with autistic features and mild motor delays while others have a progressive neurodegenerative disease (Chong *et al,* 2016). We have further characterized a cohort of Puerto Rican children sharing a *TBCK* truncating mutation (p.R126X) who exhibit the severe neurodegenerative phenotype with distinctive motor neuron degeneration, which we called TBCK-Encephaloneuronopathy (TBCKE) (Ortiz-Gonzalez *et al,* 2018). How loss of function of TBCK protein leads to disease remains unclear, but patients exhibit clinical features reminiscent of lysosomal storage disorders, including progressively coarse facies and macroglossia. Interestingly, other symptoms overlap with mitochondrial disease, such as metabolic strokes and neurologic decompensations with febrile illness.

*TBCK* encodes a protein of unclear function, <u>TBC1</u>-domain containing <u>K</u>inase. It consists of a TBC (Tre-2, Bub2, and Cdc16) domain flanked by an N-terminal kinase-like domain and a rhodanese homology domain at the C-terminus. TBCK mRNA and protein appear to be expressed in most tissues of the human body (Uhlen *et al,* 2015). Mouse brain transcriptome data shows that TBCK is expressed in astrocytes, neurons and oligodendrocytes (Zhang *et al,* 2014). Based on sequence homology with other TBC1-domain proteins, TBCK is predicted to function as a Rab GTPase-activating protein. Rab GTPases are key regulators of membrane trafficking (Kiral *et al*, 2018). Defects in Rab proteins and/or Rab GTPases leading to altered membrane trafficking have also been linked to common neurodegenerative disorders (Kiral *et al.*, 2018; Shi *et al,* 2017). These proteins are thought to be particularly critical for neuronal health because trafficking and maturation of endosomes, autophagosomes and lysosomes is crucial for successful degradation of senescent proteins and organelles (Ao *et al,* 2014; Bento *et al,* 2013)

Furthermore, a critical developmental role of TBC1-domain containing family of Rab GTPases is recently emerging. In addition to TBCK, loss of function variants in genes encoding some of these proteins are associated with neurodevelopmental and intellectual disability syndromes, such as TBC1D24, TBC1D7, TBC1D20 and TBC1D23.

Before *TBCK* was associated with human disease, studies had suggested that *TBCK* knockdown lead to downregulation of mTOR (mechanistic target of rapamycin) signaling in various cellular models (Liu *et al,* 2013; Wu *et al,* 2014). The mTOR pathway regulates crucial cellular responses including growth, apoptosis, autophagy, translation, energy metabolism, and inflammation (Laplante & Sabatini, 2012). The mTOR protein is a kinase that interacts with several proteins to form two distinct complexes named mTOR complex 1 (mTORC1) and mTOR complex 2 (mTORC2). Patient-derived TBCK-deficient cells showed a 78% decrease in phospho-ribosomal S6 phosphorylation (PS6), a widely used marker for mTORC1 downstream signaling. Moreover, the phosphorylation defect of S6 in patient cells could be rescued by L-leucine, an mTORC1 signaling activator (Bhoj *et al,* 2016). Therefore, understanding the downstream consequences of mTOR signaling inhibition in TBCK syndrome could be not only relevant in informing disease mechanisms, but also could inform potential therapeutic targets.

The mTOR pathway is known to regulate autophagy, the physiologic process by which proteins, lipids and organelles are trafficked to lysosomes for degradation (Menzies *et al.*, 2015; Tsukada & Ohsumi, 1993). We first examined autophagic function in TBCKE (ie severely affected neurodegenerative phenotype) and found increased autophagic flux and impaired degradation of glycosylated proteins in TBCKE patient derived fibroblasts (Ortiz-Gonzalez *et al.*, 2018). The impairment of autophagic-lysosomal function, was also evident in aberrant oligosaccharide degradation profiles in TBCKE fibroblasts and was ameliorated by treating cells with L-leucine. Concurrently, abnormal oligosaccharide profiles were detectable in the urine of TBCKE patients, suggesting that impaired degradation of glycosylated proteins could provide a noninvasive biomarker of impaired autophagic-lysosomal degradation in TBCK syndrome (Ortiz-Gonzalez *et al.*, 2018).

Given these previous data the first goal of the present study was to examine if the autophagic degradation of mitochondria (mitophagy) is altered in TBCK. Defects in mitophagy have been linked to neurodegenerative disorders, most prominently Parkinson’s disease (Narendra *et al,* 2008, 2009). Mitophagy is induced by mTORC1 inhibition (Pan *et al,* 2009) as well as by disruption of lysosomal acidification (Ding & Yin, 2012), both of which may play a role in TBCKIn the present study we examined mitochondrial function as well as mitophagy in TBCK encephalopathy. In particular, we were interested in examining if mitochondrial function varies in patient of diverging clinical severity and could predict a neurodegenerative course. Our data suggests, for the first time in TBCK syndrome, that indeed secondary mitochondrial dysfunction predicts a neurodegenerative course in this disease.

Finally, since effective autophagy is ultimately dependent on successful lysosomal degradation of substrates, and recent neuropathological reports suggest a lysosomal storage-like phenotype in TBCK patients (Beck-Wodl *et al,* 2018; Sumathipala *et al,* 2019), we assayed lysosomal function in TBCK patient cells.

Our data suggest that lysosomal dysfunction may be a final common pathway that could explain a blockage in degradation of mitophagosomes as well as drive mTORC1 signaling inhibition in TBCKE.

## Materials and Methods

### Cell culture

Primary patient fibroblasts were obtained with appropriate consent under an IRB-approved protocol at the Children’s Hospital of Philadelphia. Fibroblasts were maintained in DMEM with glutamax media supplemented with 10-15% fetal bovine serum (FBS) and non essential amino acids (NEAA). Experiments were carried out with cells of comparable passage number, and cells were not used past passage 15. Fibroblast lines derived from patients homozygous for the Boricua mutation (p.R126X), with a clinical course of progressive encephalopathy and motor neuronopathy are defined as “severe” (n=4 individual lines from unrelated patients, 126-1, 126-2, 126-3, 126-5). None of the p.R126X patients achieved language or independent ambulation. The “mild” phenotype lines are derived from 3 patients with no clinical evidence of neurodegeneration. Two of the patients are siblings with global developmental delay but achieving expressive language and independently ambulation (although delayed, around age 2-3yo, previously reported by Bhoj, et al.). “Mild” patients genotypes are as follows: lines 2685 and 2686 are compound heterozygous for a splice site and a frameshift variant (c.[2060-2A>G]; [803_806delTGAA],p.[=];[Met268fsArg*26] and line 16-3607 is homozygous for exon 21 deletion. Control fibroblast cell lines (similar age individuals) were obtained from the NIGMS Human Genetic Cell Repository at the Coriell Institute for Medical Research: GM17064, GM17071 and GM08398B.

### Mitochondrial Assays

Mitophagy was assayed by live confocal imaging of primary TBCKE fibroblasts at basal culture conditions. Cells were plated in glass bottom 35 mm dishes (MatTek) and incubated with Mitotracker Green (thermofisher) 20 nM for 15 minutes and Lysotracker Red (thermofisher) 50 nM for 25 minutes, and washed with pre-warmed PBS. Cells were imaged in a LSM710 (Carl Zeiss) confocal microscope equipped with temperature controlled stage. Alternatively to assay mitophagy by genetically encoded labels, fibroblasts were treated simultaneously with the LC3B-GFP (premo atutophagy sensor Thermofisher P36235) and mito-RFP (thermofisher C10601) per manufacturer’s inctructions. Colocalization index was determined using Zeiss Zen blue software using the same acquisition and analysis settings for all images.

Mitochondrial Respiration: Baseline respiration and maximal respiratory capacity were measured using the Seahorse instrument (XF24 data for Figure 2 and Xe96 data for figure 3) according to manufacturer’s instructions (Agilent). Briefly, 20,000 cells were seeded per Xe96 plate well and cells were assayed within 24 hours of plating unless otherwise noted. The following reagent concentrations were used: Oligomycin (1.25uM), FCCP (1 uM), antimycin (1.8uM) and rotenone (1uM). For respiration assays with L-leucine treatment: Cells were plated as above but supplemented for 24 hours with 600 ug/mL of L-leucine, as this dose was previously reported to augment mTORC1 mediated phosphorylation in TBCK cells (Bhoj *et al.*, 2016).

### Mitochondrial Content

Mitochondrial mass was evaluated using anti-VDAC-1 staining quantified by flow cytometry. Briefly, cells were collected and washed with cold phosphate buffer saline (PBS), fixed with 4% PFH, and kept on FACS buffer (PBS, 0.5% BSA, 0.05% Sodium Azide). For staining cells were washed twice with Intracellular Staining Permeabilization Wash Buffer (Biolegend catalog No. 421002), and re-suspended in same buffer with anti-VDAC-1 (abcam catalog No.15895) for 30 min at room temperature. Cell samples were washed with wash buffer and and incubated secondary (Alexa Fluor® 647Anti-Rabbit IgG Jackson Immuno Research Laboratories catalog No.111-605-003) for 30 at room temperature in dark. After two washes with FACS buffer cells were re-suspended on FACS buffer and assessed on CytoFLEX flow cytometer (Beckman Coulter, Life Science, Indianapolis, IN, US). Flow data analysis was performed using FlowJo software.

### Mitochondrial DNA copy number and Gene expression Assays

mtDNA copy number was assayed in fibroblasts from patients with TBCK mutations but distinct clinical phenotypes as described above. “Mild” patients without evidence of disease progression (n=3 patient lines) were compared with the “severe” Boricua p.R126X mutation (n=4 patient lines). Control lines were acquired commercially as detailed above. For mtDNA copy number DNA was isolated using manufacturer’s protocol DNeasy Blood and tissue kit (QIAGEN Catalog No. 69506). Published primer sequences for CytB, ND4, ND1, tRNA Leu (UUR), HRPT, B2M, 18s were used (Kaaman *et al,* 2007; Venegas & Halberg, 2012)

For gene expression experiments, extraction of total RNA was performed using Rneasy Plus Universal mini kit (QIAGEN Catalog No. 73404), total RNA samples were processed with TURBO DNA-free^™^ Kit (ThermoFisher Scientific Inc., Catalog No. AM1907) (1 U/1 μg RNA), for 30 min at 37 °C, in order to avoid contaminant genomic DNA. Then, reverse transcription was performed using the SuperScript™ IV Reverse Transcriptase kit (ThermoFisher, Catalog No. 18090010), with Oligo(dT)20 primer. For each cDNA sample, qPCR was performed with the following amplification primers: PGC1a (5’-TCC TCT TCA AGA TCC TGC TAT T3’; antisense, 5’ ACG GCT GTA GGG CGA TCT 3’). Stably expressed reference gene, GAPDH (sense, 5’ TCG GAG TCA ACG GAT TTG G 3’; antisense, 5’ TCG CCC CAC TTG ATT TTG GA 3’), was used for target gene expression normalization.

Amplifications were carried out in 384-well plates on VIIA7 thermocyler and each sample was analyzed in triplicate. Negative controls without template were run for each gene. The reaction solution was prepared by combining 30 ng of DNA to each well as the template, 1X SYBR Green PCR Master Mix (Applied Bio systems) and 0.4 mM of each of primer for a total volume of 10 μl. Amplification program: 10 min at 95 °C, followed by 40 cycles of 15 s at 95 °C and 60 s at 60 °C. At the end of the amplification process, the melting curve was performed to assess amplification specificity of each gene. The corresponding real-time PCR efficiencies for each mitochondrial and nuclear gene amplification were calculated according to the equation: E=10(−1/slope)−1. Relative mtDNA copy number (mtDNA amount/nDNA amount) and relative expression levels of PGC1α was calculated by a comparative Ct method

### Lysosomal Assays

Cathepsin D: Freshly isolated fibroblasts lysates were used to measure cathepsin D activity using a fluorometric plate reader assay per manufacturer instructions (abcam ab65302).

DQ-BSA: Fibroblasts were incubated for 4 hours at 37C with 10ug/ml DQ™ Green BSA (ThermoFisher Catalog No. D12050). Prior to staining cells were serum deprived for 4 hours to maximize uptake. To further confirm live imaging findings and quantify proteolytic degradation, cells were also assayed by flow cytometry. Fibroblasts were treated with DQ-BSA vs blank controls and mean fluorescence intensity (FITC) was quantified using flow cytometry (CytoFLEX) and data analyzed with FlowJo.

Neutral lipid staining was done with live fibroblasts treated with Bodipy D3922 1μm for 30 minutes prior to live confocal imaging.

Filipin staining: Fibroblasts were washed twice in PBS, fixed with 3% paraformaldehyde for 30 min at room temperature (RT) followed by 2 washes with PBS and incubated in quenching solution (1.5 mg/ml glycine, 1X PBS) at RT for 10 min. This was followed by incubating cells in 0.5 mg/mL filipin (ThermoFisher, Catalog No. D3922), 10% FBS for 2 hours at RT in the dark. After washing with PBS cells were imaged using a Zeiss LSM 710 confocal microscope.

### Nanoparticles

Acidic nanoparticles (NP) that have been previously shown effective in acidifying lysosomal pH and improving lysosomal function (Baltazar *et al,* 2012; Lee *et al,* 2015)) were a generous gift of Dr Claire Mitchell. Appropriate uptake into lysosomes in TBCK fibroblasts was verified by using Nile Red labeled NP and co-labeling with 50nM lysotracker green (figure 5). NP were resuspended in culture media (1mg/mL), then sonicated for 10 min and filtered through a 0.8 μm filter (Corning, Catalog No. 431221). Fibroblasts were incubated in NP media for 3 days and then mitochondrial respiration was assayed using seahorse instrument as described above.

All statistical analysis was performed using Graph Pad Prism, with p value <0.05 considered significant. For all figures, * p<0.05; ** p<0.01, *** p<0.001, **** p<0.0001

## Results

### TBCKE fibroblasts exhibit upregulation of mitophagy and reduced mtDNA content

Since we previously reported increased autophagic flux in TBCKE fibroblasts (Ortiz-Gonzalez *et al.*, 2018), we first tested whether mitophagy was altered in TBCK patient derived versus control fibroblasts. Live confocal imaging shows extensive colocalization of functional mitochondrial and lysosomal markers, an indicator of mitophagy. Figure 1a shows representative images and the quantification of the colocalization coefficient in control vs TBCKE fibroblasts at baseline culture conditions (n=2 control lines and 2 TBCKE patient lines homozygous for p.R126X mutation). In order to confirm the upregulation of mitophagy with a different technique, we transduced control and TBCK fibroblasts with mito-RFP and LC3-GFP. We observed mitochondria clearly enclosed in LC3-GFP positive autophagosomes in TBCK fibroblasts, and confirmed significant increase in colocalization coefficient relative to control (Extended view figure 1)

**Figure 1:**
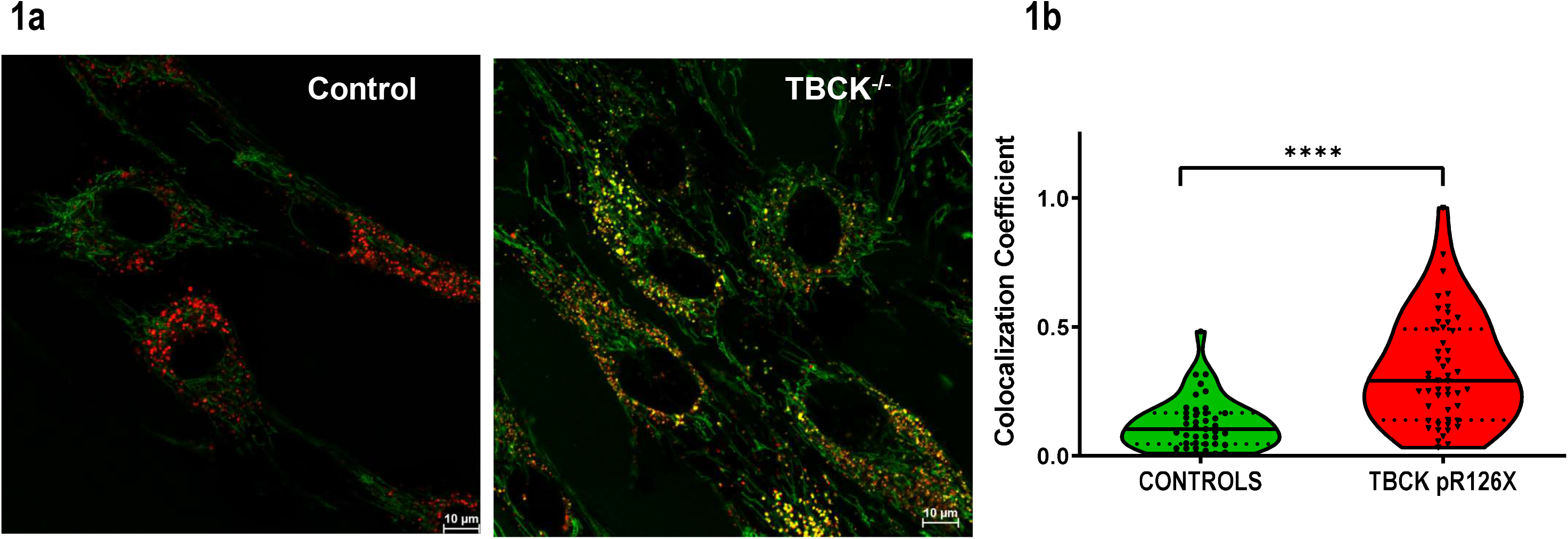
Mitophagy is upregulated in TBCKE fibroblasts. Panel (a) shows confocal live imaging of human fibroblasts at baseline culture conditions, stained with mitotracker green and Lysotracker red. TBCKE (p.R126X) fibroblast exhibit robust colocalization, a marker of mitophagy. Panel (b) shows quantification of colocalization index in 4 individual cell lines (2 controls, circles and 2 TBCK p.R126X lines, triangles, 126-2 and 126-3), p<.0001, One way Anova

**Figure 2:**
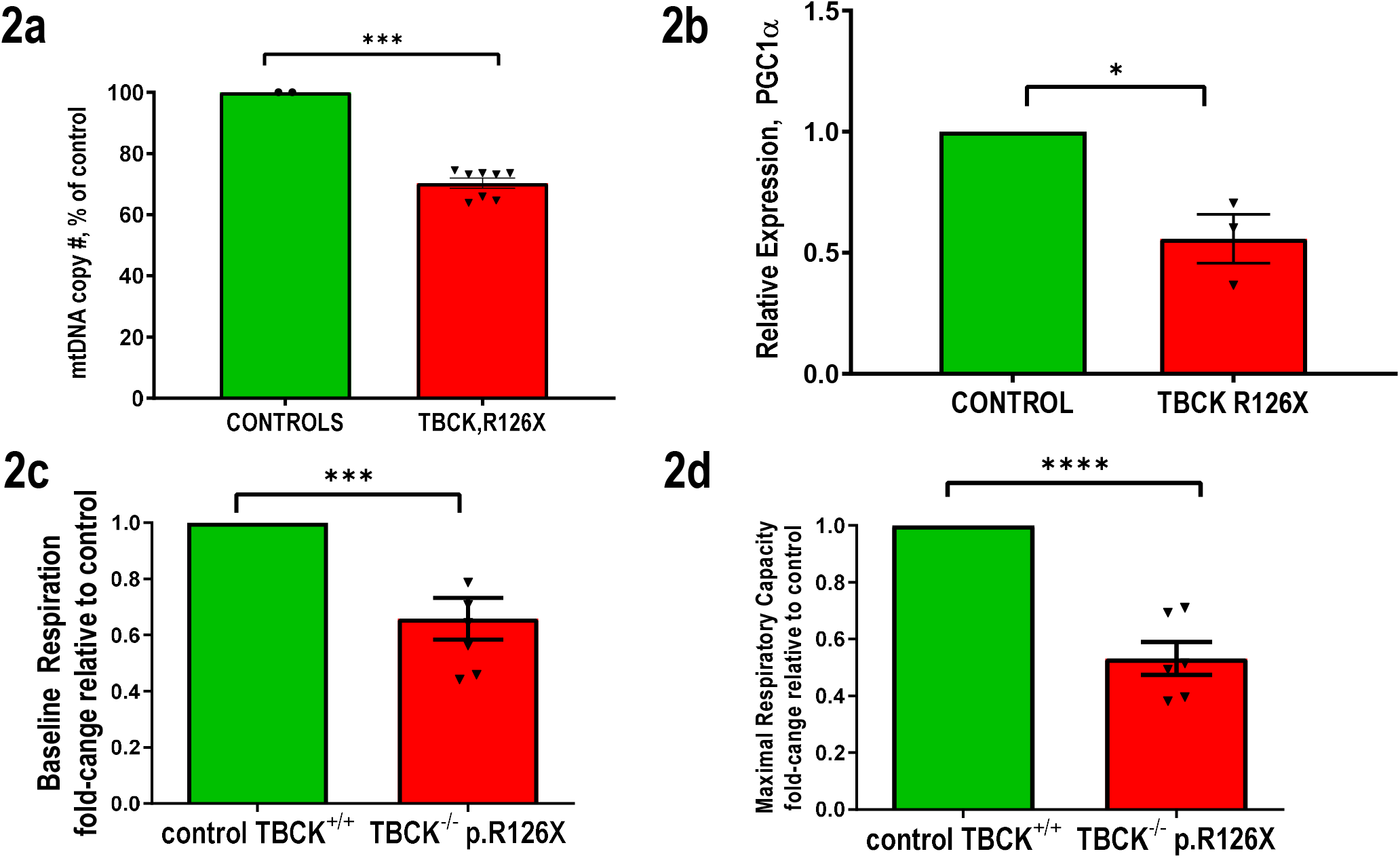
TBCKE fibroblasts have reduced mitochondrial DNA and respiratory function. (2a) mtDNA copy number in 3 independent patient-derived lines homozygous for the TBCK Boricua (p.R126X, 126-1, 126-3 AND 126-5) mutation show 30% reduction in mtDNA, normalized to controls (mean 70.3%, P<.0001 unpaired t-test). (2b) Expression of PGC1α, a key modulator of mitochondrial biogenesis is also decreased in TBCK patient cells (mean .55, p value .0116, unpaired t-test). Seahorse assays (n≥4 assays) of TBCK fibroblasts (p.R126X) normalized to controls show (2c) reduced baseline mitochondrial respiration (66% of control, p .007 unpaired t-test) as well as (2d) reduced FCCP stimulated maximal respiratory capacity (53% of controls, p <.0001 unpaired t-test).

Since we found evidence for upregulation of mitophagy, we then asked if mtDNA copy number was reduced in TBCKE fibroblasts, as mtDNA damage is known to trigger mitophagy (Gilkerson *et al,* 2012). Figure 1b shows that mtDNA copy number is significantly reduced in TBCK fibroblasts from Boricua mutation patients. mtDNA copy number is controlled by the interplay between mitochondrial biogenesis and degradation via mitophagy. In order to gain insight into whether less mtDNA in TBCK cells could be secondary to decreased generation of mitochondria, we examined the expression of *PGC1α*, a transcription of co-activator that regulates mitochondrial biogenesis. We found that *PGC1 α* expression was also significantly reduced in all TBCKE patient lines (figure 1c).

Given increased mitophagy and reduced mtDNA content, we then tested mitochondrial function by measuring respiratory capacity in TBCKE fibroblasts. Figure 1b shows that both baseline and maximal respiratory capacity are significantly reduced in TBCKE fibroblasts (with Boricua mutation p.R126X). Baseline respiratory capacity was 66% of controls, while maximal, FCCP stimulated respiratory capacity was 53% of control levels.

### Mitochondrial dysfunction correlates with severity of clinical phenotype

Mitochondrial respiration was measured in cell lines derived from patients with evidence of neurodegeneration (“severe”, n=4 lines, all with the Boricua p.R126X homozygous mutation) and compared to cell lines from patients with a static clinical course (“mild”, n=3 lines, see methods sections for genotypes). Although all *TBCK*^-/-^ patient lines exhibited significant mitochondrial respiration defects relative to controls (figures 3a–3b), fibroblasts from severe patients had significantly lower maximal respiratory capacity than cells from mild patients (figure 3c). Interestingly, at baseline, mild vs severe phenotypes have similarly reduced oxygen consumption rates (3b). This suggests that the severity of mitochondrial respiratory defects, specifically, the spare respiratory capacity, correlates with the severity of neurological deficits in patients with *TBCK* mutations.

**Figure 3:**
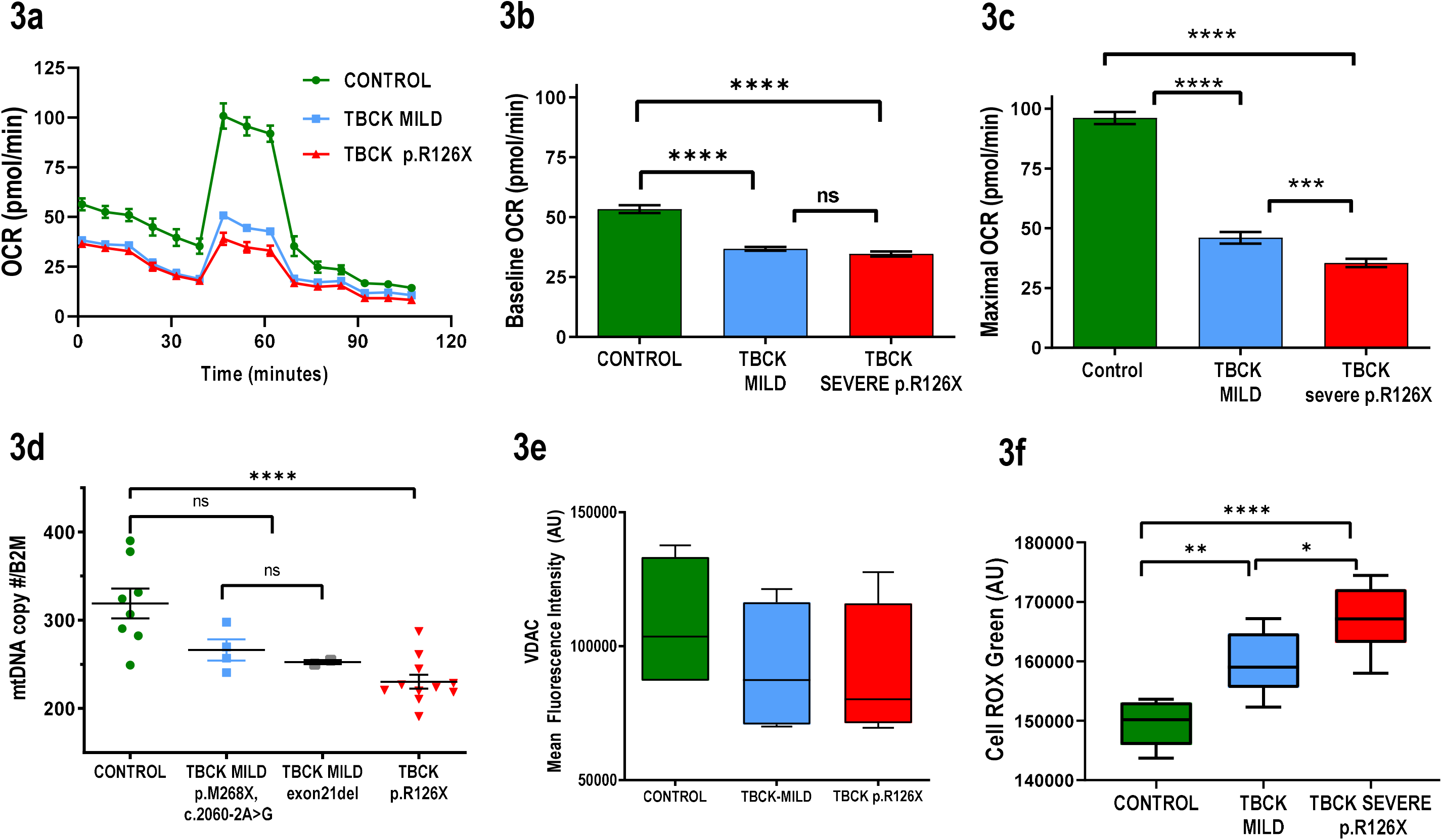
Reduced mitochondrial respiratory capacity and mtDNA correlate with severity of neurologic phenotype in TBCKE. Mitochondrial oxygen consumption rate (OCR) of control compared to *TBCK*^-/-^ patient lines with mild (blue) vs severe (red) neurologic phenotypes. Both “mild” and “severe” groups included fibroblasts lines derived from at least 3 individual patients (see methods for details and genotypes). Panel a shows representative OCR tracing plotted in panel 3b for baseline respiration and 3c for maximal respiratory capacity, showing that defects in maximal respiratory capacity predicts the severity of the clinical phenotype. Similarly, mtDNA content (3d), (normalized to nuclear gene *B2M*) correlates with severity of neurologic phenotype, with mild TBCK patients exhibiting similar mtDNA levels as controls while severe Boricua patients (p.R126X) have significantly decreased mtDNA content. Mitochondrial content per cell, assayed by mitochondrial outer membrane protein porin (VDAC) staining quantifed by flow cytometry, does not significantly differ in control vs TBCK lines (3e), while levels of reactive oxygen species (ROS) do vary significantly with genotype (3f).

To further examine the correlation of mitochondrial measures with phenotype, we measured mtDNA copy number in fibroblasts from severe vs mild patients, relative to healthy controls. Consistent with our respirometry data, mtDNA copy number was reduced in all TBCK patient lines, but most strikingly in the fibroblasts derived from the severely affected TBCKE patients with the Boricua mutation (mild 46% of control, severe 36% of control, Figure 3d).

### Reactive Oxygen Species (ROS) levels, but not mitochondrial content, are significantly altered in TBCK cells

In order to understand why mtDNA content may be reduced in TBCK cells, we first examined whether there is reduced amount of mitochondria per cell. As a measure of mitochondrial content, we quantified the amount of VDAC-1 (porin), one of the most abundant outer membrane mitochondrial proteins which is encoded by nuclear DNA (Janes *et al*, 2004). Multiple experiments quantifying porin by western blot and flow cytometry did not show a consistent decrease in mitochondrial content in TBCK cells. Figure 3e shows flow cytometry quantification, despite a trend to lower mitochondrial content in TBCK affected cells, there was no statistically significant reduction in mitochondrial content per cell. Citrate synthase assays, another measure of mitochondrial content, also showed no significant differences between TBCK^-/-^ and control lines (data not shown). If mtDNA copy number reduction is not due to reduced amount of mitochondria per cell, an alternative scenario is reduced amount of mtDNA per mitochondria. Reactive Oxygen Species (ROS) generated in the mitochondria are known to induce mtDNA damage(Nissanka & Moraes, 2018). Therefore, we next assayed ROS levels at baseline conditions using CellROX in controls vs TCBK (mild and severe) fibroblasts. We found a striking increase of ROS levels in TBCK cells, and furthermore, significant correlation with clinical severity (figure 3f). This observation is consistent with our mtDNA content data, and implicates that, concomitant with reduced PGC1α expression, elevated ROS levels may contribute to mtDNA depletion in TBCK.

### TBCK fibroblasts have reduced lysosomal proteolytic activity

TBCK^-/-^ fibroblasts had a moderate but significant decrease in enzymatic activity of the lysosomal protease cathepsin D (27% reduction, n=2 control lines and n=4 severe lines, Fig 4a). To further examine lysosomal proteolytic function in fibroblasts, we used DQ-BSA. Fluorescence of this molecule is quenched at baseline, but activated once proteolytic degradation occurs within the acidic lysosome. We found by confocal imaging that DQ-BSA signal was visibly reduced in TBCKE fibroblasts (fig 4b), which was confirmed and quantified using flow cytometry (Fig 4c). Fibroblasts derived from severe TBCKE Boricua patients had a significant 35% reduction in DQ-BSA proteolytic degradation relative to controls (n=2 cell lines per group). Furthermore, we observed the same trend as we saw when quantifying mitochondrial function; that is, a significant impairment in lysosomal proteolytic activity correlates with the severity of clinical phenotype. We also observed accumulation of lipid droplets and unesterified cholesterol (by filipin staining) in TBCK cells, all features consistent with cellular lysosomal dysfunction (figure 4d)

**Figure 4:**
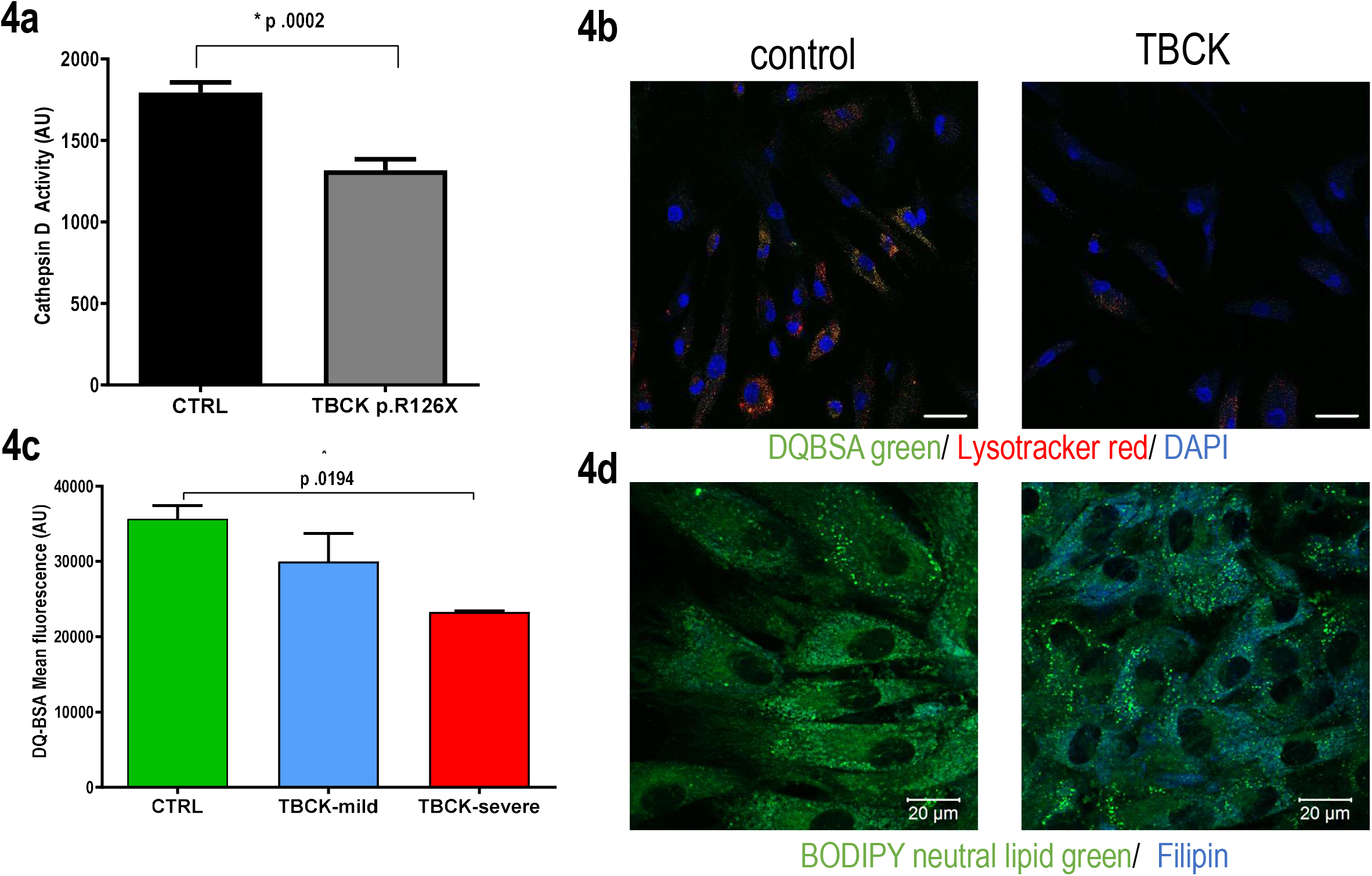
Lysosomal Dysfunction in TBCKE fibroblasts. Lysosomal proteolytic activity assays: (4a) shows reduced enzyme activity of cathepsin D and panels (4b) show reduced protelolytic processing of DQ-BSA by live imaging and (4c) flow cytometry. DQ-BSA fluorescence is dependent on lysosomal degradation. Neutral lipid staining (BODIPY493) showing increased lipid droplets (4d, green); filipin stain (4d blue) also shows unesterified cholesterol accumulation in TBCKE fibroblasts. Scale bar = 20 μm

### Acidifying lysosomal nanoparticles, but not mTOR activation with L-leucine, rescue mitochondrial respiratory dysfunction in TBCK fibroblasts

In order to test if the mitochondrial dysfunction observed in TBCK could be secondary to lysosomal pathology, we tested whether acidifying nanoparticles that are incorporated into the lysosomes (Baltazar *et al.*, 2012) rescue the mitochondrial respiratory defects in TBCK cells. TBCK and control fibroblasts were treated for 3 days with acidifying nanoparticles and mitochondrial respiration studies performed as previously. We found that all TBCK fibroblasts lines improved their mitochondrial maximal respiratory capacity after nanoparticle treatment (Figure 5a). Figure 5B shows colocalization of nile-red labeled nanoparticles with lysotracker green, in order to confirm appropriate trafficking of nanoparticles into the lysosomes. On the other hand, L-leucine treatment, which has been shown to positively modulate mTORC1 mediated phosphorylation in TBCK cells (Bhoj *et al.*, 2016) did not alter the mitochondrial respiratory defects (supplemental figure 2).

**Figure 5:**
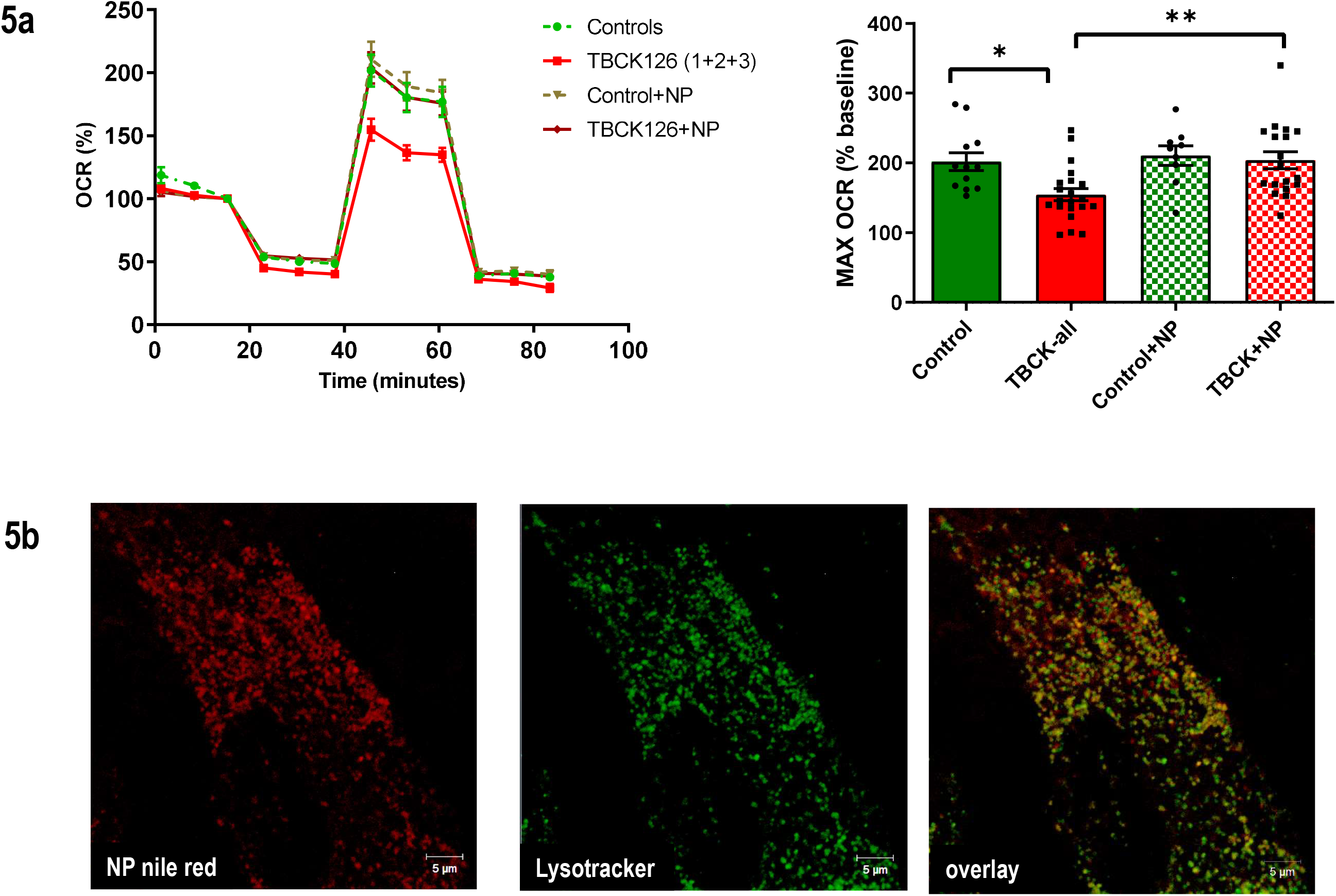
Lysosomal Acidification rescues mitochondrial respiratory defects in TBCKE. fibroblasts. Control and TBCK fibroblasts were treated for 3 days with acidifying nanoparticles (courtesy of Dr Claire Mitchell) and mitochondrial respiration assayed as previously using seahorse XF96 instrument. Control lines (n=2) showed no change in respiratory capacity after treatment, while mitochondrial respiratory defects in TBCK fibroblasts (n= 3 lines) treated with NP were restored to control levels. Panel B shows control fibroblasts treated with nile-red nanoparticles colocalized with lysotracker green, verifying that NP are incorporated into the lysosomes (scale bar 5 μm).

## Discussion

Although we have made great advances in identifying novel disease genes in the next generation sequencing era, understanding how genetic changes affect biologic function in order to provide more accurate prognosis and ultimately, treatments, remain a major challenge in the neurogenetics clinic. This is particularly challenging for novel disease genes with highly variable neurologic outcomes, such as the neurodevelopmental syndrome associated with *TBCK* mutations. Most, if not all, patients share significant hypotonia and motor delays in infancy and early childhood. Some patients proceed to gain developmental skills such as independent ambulation and language, and to date have not shown evidence of regression. Most commonly though, children experience progressive neurologic decline with clear brain atrophy on serial imaging and evidence of motor neuron degeneration, which ultimately can be fatal due to chronic respiratory insufficiency. Genotype-phenotype correlations are not completely clear cut, severely affected patients usually have truncating mutations early in the protein, but some of the mild patients also have splice site mutations that prevent protein expression and lead to undetectable protein levels (Bhoj *et al.*, 2016). Our group previously predicted, using *in silico* modeling, that most disease causing variants could alter the GAP activity domain (Chong *et al.*, 2016). In the present study, we examined primary fibroblasts from patients with biallelic mutations in *TBCK*, and assayed for biological measures that may give us mechanistic insight into its pathophysiology, as well as inform what cellular functions correlate with disease severity.

The physiologic function of TBCK remains unclear. It is predicted to act as a RabGTPase, based on sequence homology. Unbiased screening methods to detect effectors of Rab GTPases have found TBCK to bind to Rab5 in *drosophila* (Gillingham *et al*, 2019; Gillingham *et al*, 2014) but to our knowledge there is no clear published data to confirm the TBCK function or binding partners of TBCK in mammalian cells.

Other TBC1-domain containing proteins have also been proposed to interact with Rab 7 and play a role in regulating delivery of clathrin-coated vesicles to the lysosome (Frasa *et al*, 2012). Rab7a is known to regulate endosomal transport and late endosome-lysosome fusion, but recent data specifically suggest a role in regulating autophagosome biogenesis during mitophagy, also known as mitophagsome formation (Tan & Tang, 2019; Yamano *et al*, 2014). It has been shown that Rab7A phosphorylation via TBK1 promotes mitophagy via the pink-parkin pathway (Heo *et al*, 2018). Rab7 GTP hydrolysis has also been implicated in regulating mitochondria-lysosomal contact sites, controlling mitochondrial fission (Wong *et al*, 2018).

Since autophagic-lysosomal clearance is abnormal in severe TBCKE (Boricua mutation) cells, the present study focuses on mitophagy specifically, the physiologic removal of mitochondria by the autophagic cellular machinery. Previously, we showed that TBCKE cells have increased autophagic flux and impaired degradation of glycoproteins, both consistent with autophagic-lysosomal dysfunction (Ortiz-Gonzalez *et al.*, 2018). Autophagic degradation of abnormal mitochondria via mitophagy is particularly relevant to postmitotic cells like neurons. In fact, some of the first insights into the relevance of mitophagy for neurologic disease were derived from patients with Parkinson’s patients who were found to have mutations in PINK1 and parkin (PARK2) (Kitada *et al*, 1998; Matsumine *et al*, 1997; Valente *et al*, 2004a; Valente *et al*, 2004b). These studies have greatly expanded our knowledge of the role of mitochondrial quality control in various neurodegenerative syndromes, both rare and common (Audano *et al*, 2018).

Using patient derived fibroblasts, we found robust colocalization of lysosomes and mitochondria in TBCK cells at baseline culture conditions (figure 1). Therefore, we then questioned whether excessive mitophagy could lead to depletion of mtDNA in *TBCK*^-/-^ cells and subsequent mitochondrial respiratory defects. mtDNA depletion syndromes are known to cause childhood-onset neurodegeneration (Young & Copeland, 2016), and given the variable phenotype of TBCK patients, we hypothesized that impaired mitochondrial dysfunction and mtDNA copy number may distinguish the static versus neurodegenerative clinical course.

Our data shows that at baseline, both mild and severe TBCK cell lines have impaired mitochondrial respiration relative to control fibroblasts. Nevertheless, when cells were treated with an uncoupler (FCCP) to stimulate mitochondrial maximal respiration, a consistent correlation between mitochondrial respiratory capacity and severity of the phenotype emerge. Cells with a mild phenotype have preserved respiratory capacity relative to the severe lines. Consistent with this result, mtDNA copy number was lower in mild cells but not to a lesser extent relatively to the severe cell lines (see Figure 3). Therefore, control vs severe (but not mild) showed significant decrease in mtDNA copy number. We found that despite the upregulation in mitophagy markers in *TBCK*^-/-^ fibroblasts, there was no significant effect on mitochondrial content per cell in *TBCK*^-/-^ vs control cells. We did find ROS levels that also were proportional to the severity of the phenotype. Overall, this suggests that higher ROS levels and mtDNA damage may underlie the respiratory defects in TBCK fibroblasts, as opposed to a depletion of the number of mitochondria per cell. It remains to be determined whether these observations are true in neural cells, given the primarily neurologic disease phenotype.

Ultimately, autophagy, including mitophagy, relies on lysosomes for degradation of defective proteins and organelles (Menzies *et al.*, 2015; Tsukada & Ohsumi, 1993). Increased autophagic flux could be secondary to increased signaling to initiate autophagic cascades (ie via mtOR signaling inhibition), but could also be seen in association with defective lysosomal degradation. Since we reported increased autophagosomes and impaired degradation of oligosaccharides in TBCKE fibroblasts(Ortiz-Gonzalez *et al.*, 2018), neuropathological reports from deceased TBCK patients have proposed reclasifiying TBCK as a lysosomal storage disorder (LSD) based on evidence of lysosomal pathology in human brains (Beck-Wodl *et al.*, 2018). Therefore, we asked whether there is evidence of lysosomal dysfunction in TBCK patient fibroblasts, and furthermore, whether this could also be correlated with clinical phenotype.

Our data shows that there is moderate (around 30-35%) and statistically significant decrease in proteolytic lysosomal activity by both cathepsin and DQ-BSA hydrolysis assays when comparing severe *TBCK*^-/-^ patient lines to controls (Figure 5). Furthermore, the correlation with severity of phenotype is also maintained, with mild *TBCK*^-/-^ lines showing 10-15% decrease in activity that is not statistically significantly different than controls. Not only we show evidence for lysosomal proteolytic defects in TBCK fibroblasts, we also show accumulation of lipid droplets and unesterified cholesterol, reminiscent of other LSD. Therefore, our data suggests that impairment in lysosomal function leads to impaired mitochondrial quality control and subsequent mtDNA depletion and respiratory dysfunction, and that the severity of this impairment predicts the neurodegenerative course in TBCK encephaloneuronopathy.

Our data suggesting lysosomal dysfunction may also provide a mechanistic link with the previous observation of decreased mTORC1 siganaling. It has been proposed that mTORC1 activation and sensing of amino acids occurs at the lysosomal surface, and previous studies have shown that lysosomal dysfunction is sufficient to inhibit mTORC1 signaling (Bar-Peled & Sabatini, 2014; Li *et al*, 2013).

Further studies may test whether the increased number of mitophagosomes we observed could be due to TBCK leading to impaired Rab5 and/or Rab7 cycling, altering mitophagosome biogenesis and/or mitochondrial-lysosomal contacts. Our data is consistent with increased mitophagosomes being a marker for impaired lysosomal degradation. Future studies should also examine if lysosomal uptake and endosomal trafficking pathways are intact in TBCK, as this has been shown to play a role in some classic lysosomal storage disorders such as batten’s disease due to CLN3 mutations. (Schultz *et al*, 2018; Schultz *et al*, 2014).

Furthermore, we show that lysosomal rescue strategies, such as acidifying nanoparticles, effectively rescue the mitochondrial defects seen in TBCK syndrome, consistent with secondary mitochondrial dysfunction as opposed to a primary bioenergetic defect. In our studies, mTOR modulation with L-leucine did not alter the mitochondrial dysfunction. Further studies with other mTOR modulating strategies, whether genetic or pharmacological, should be considered, ideally in neuronal cell models. Future studies should also examine the value of lysosomal rescue strategies using neuronal cell models and *in vivo* using animal models to further establish their therapeutic potential for this devastating disease.

## Acknowledgements

First and foremost, we want to thank the patients and their families for their invaluable contribution to this research. We also thank our collaborators Dr. Elizabeth Bhoj and Dr. Daniel Doherty for referring patients to our cohort. Dr Claire Mitchell generously provided the nanoparticles for the lysosomal rescue studies.

This work was supported by the Robert Wood Johnson Harold Amos Faculty Development Award (XOG), the CHOP Roberts Collaborative (XOG) and the CHOP Foerderer Award.

## Author Contributions

JATH: Experimental design, data acquisition and analysis, editing manuscript. KK: assistance with human subjects recruitment, obtaining patient cell lines, editing manuscript. AS: data acquisition and analysis, editing manuscript. XOG: Design and direct project, experimental design, data acquisition and analysis, human subject recruitment, wrote manuscript

## The paper explained

### PROBLEM

Pediatric neurodegenerative syndromes are rare but devastating disorders, robbing children of developmental milestones at an early age. Biallelic variants in the *TBCK* gene cause a neurodevelopmental syndrome of peculiar variability, with some children exhibiting stable intellectual disability while others experience progressive neurodegeneration. The biological underpinnings that underli this phenotypic variability remain unclear, as does the function of TBCK protein

### RESULTS

This study shows that variants in TBCK lead to dysfunction of 2 critical and interconnected cellular organelles, mitochondira and lysosomes. We show increased mitophagosomes and mitochondrial dysfunction. The degree of mitochondrial respiraratory defects correlate with the severity of the phenotype. Furthermore, we show that the defects in mitochondrial degradation are secondary to lysosomal dysfunction, and rescuing lysosomes rescues the mitochondrial defects.

### IMPACT

Aberrant mitochondrial quality control has been widely implicated in neurodegeneration. Our study suggests that TBCK protein may play a role in autophagic-lysosomal cascades. Our data also suggests that it may be feasible to establish biomarkers to predict the severity of TBCK phenotype based on measures of mitochondrial and lysosomal function. Our data suggests that lysosomal acidification strategies may have therapeutic potential in TBCK encephaloneuronopathy.

## Abbreviations

mtDNA: mitochondrial DNA
TBCK: TBC1-domain containing kinase
TBCKE: TBCK Encephaloneuronopathy

## Expanded View Figure Legends

**Expanded View Figure 1:**
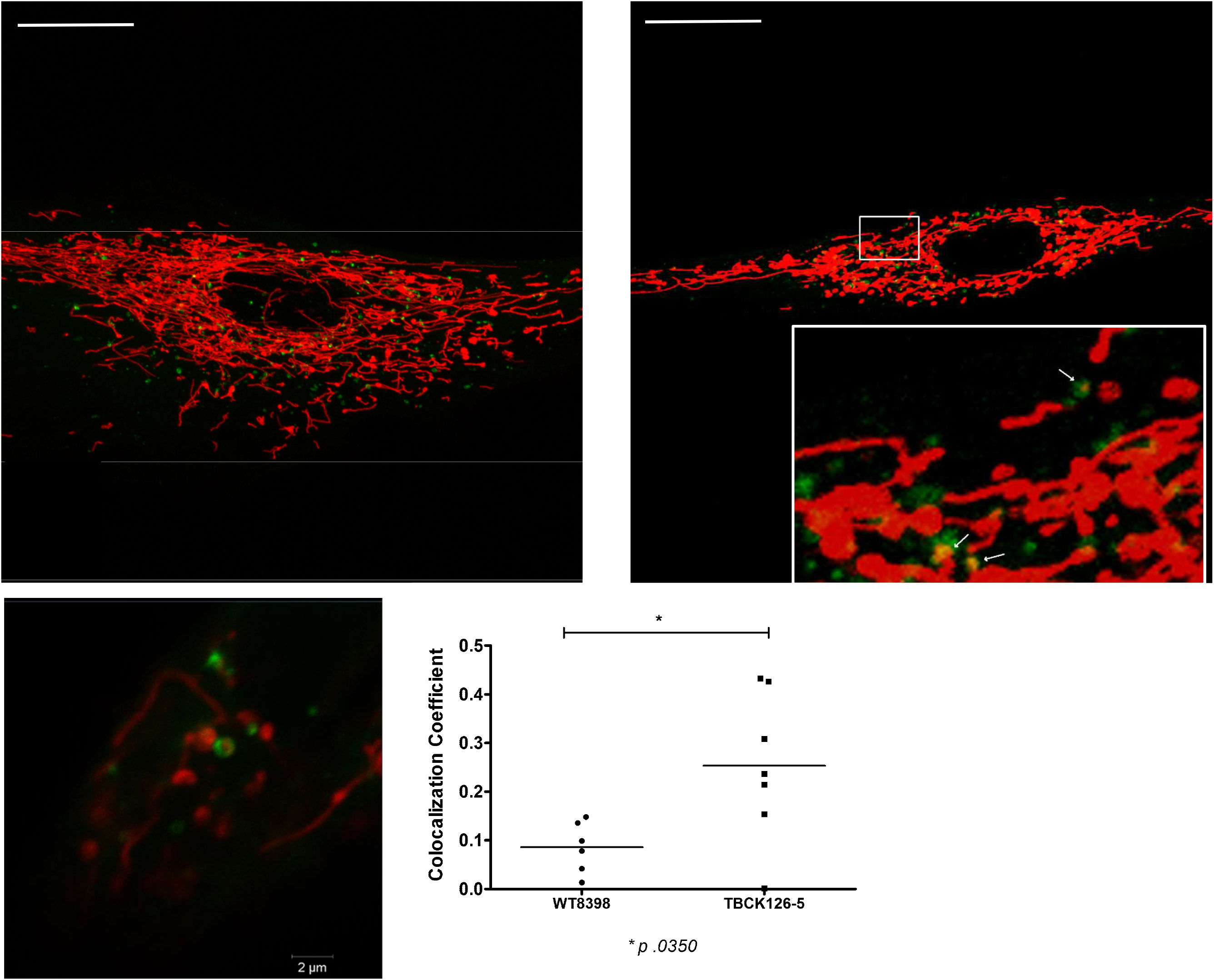
Mito-RFP and GFP-LC3 expression confirms mitochondria enclosed within autophagosomes are significantly increased in TBCKE fibroblasts. Control and TBCK fibroblast were cotransfected with mito-RFP (red) and LC3-GFP (green), live confocal imaging and colocaization analysis was performed as previously showin if fugre figure 1. Therefore, confirming with alternative labeling methods the observation of increased mitophagosome content in TBCK cells

**Expanded View Figure 2:**
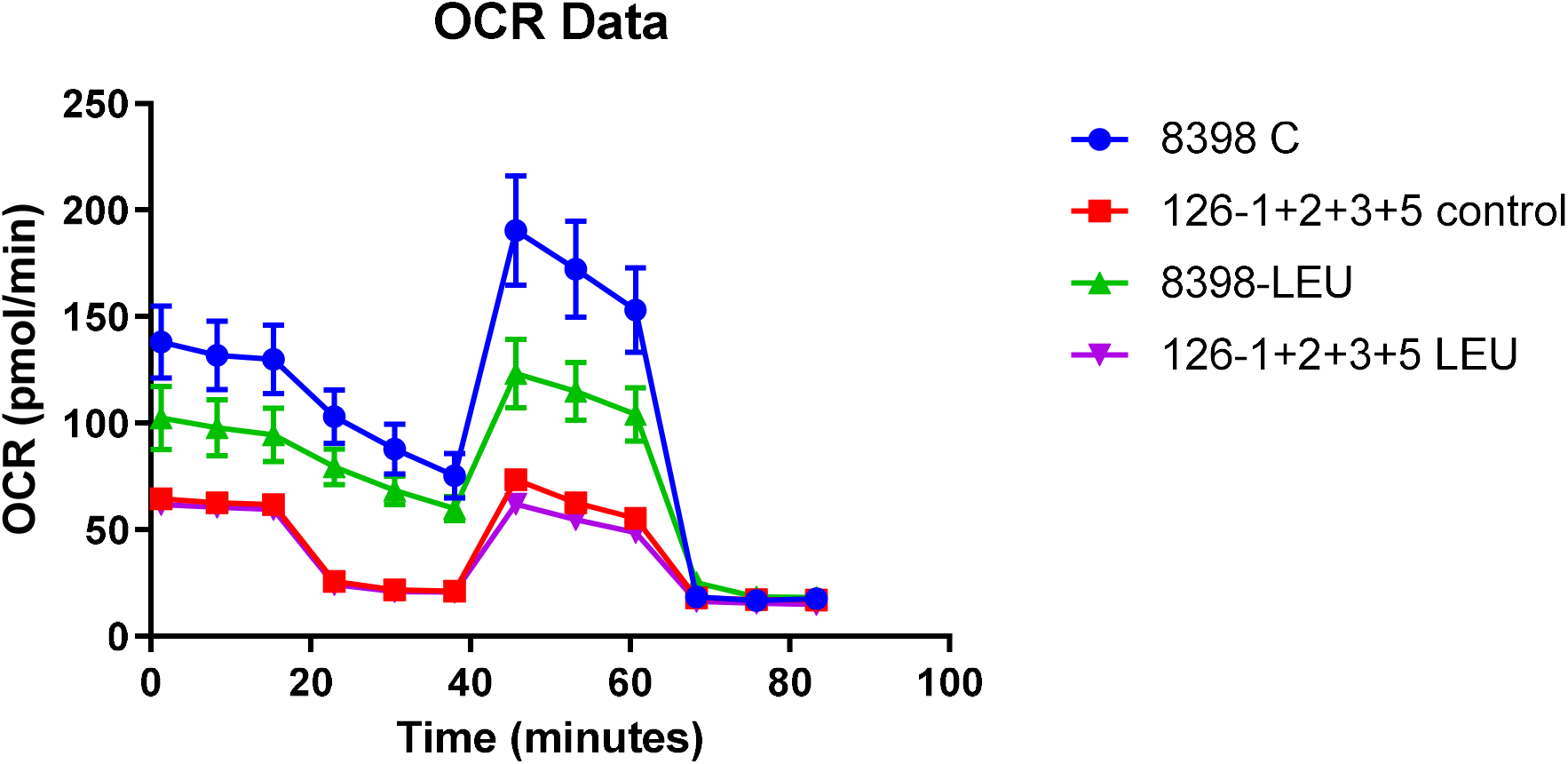
L-Leucine treatment (600 ug/ml for 24 hrs) does not rescue mitochondrial respiratory defects in TBCKE fibroblasts. Representative seahorse assay to determine oxygen consumption rate (OCR); 72 hrs of L-leucine treatment (data not shown) yielded similar results. CONTROL =8398; TBCK= 126-1, 126-2, 126-3, 126-5

